# Diverging transposon activity among polar bear sub-populations inhabiting different climate zones

**DOI:** 10.1101/2024.12.04.626794

**Authors:** Alice M. Godden, Benjamin T. Rix, Simone Immler

## Abstract

A new subpopulation of polar bears (*Ursus maritimus*) was recently discovered in the South-East of Greenland. This isolated colony inhabits a warmer climate zone, akin to the predicted future environments of polar bears with vastly reduced sea ice habitats, rendering this population of bears particularly important. Over two-thirds of polar bears will be extinct by 2050 with total extinction predicted by the end of this century, therefore understanding possible mechanisms of adaptation via genomic analyses and preservation are critical. Transposable elements (TEs) are parasitic mobile elements that may play a role in an adaptive response to environmental challenges. We analysed transcriptome data from polar bear sub-populations in cooler North- East (NEG) and warmer South-East Greenland (SEG) to compare TE activity between the two populations and its correlation with temperature and associated changes in gene expression. We identified activity hotspots in the genome of regions with significantly differentially expressed TEs. LINE family TEs were the most abundant, and most differentially expressed and divergent in the SEG population compared to reference TEs. We report a significant shift in TE activity and age, with younger more abundant TEs in the SEG populations. Differentially expressed genes in SEG populations were linked to *Foxo* signalling, ageing and metabolic pathways. Our results provide insights into how a genomic response at the TE level may allow the SEG subpopulations to adapt and survive to climate change and provides a useful resource for conservation in polar bears.

## Background

Global average temperatures in 2024 were more than 1°C above the pre-industrial era ^1^. The Arctic current is the warmest it has been in the last 125,000 years and temperatures continue to rise ^2,3^. In the South East of Greenland (SEG), the ice sheet margin is rapidly receding, causing vast ice and habitat loss ^4^. North-East Greenland (NEG) is an Arctic tundra, whereas SEG is covered by forest tundra ^5^. The SEG climate has high levels of precipitation, wind, and steep coastal mountains ^6^. This challenging habitat has led to severe warnings for the survival of the polar bear *Ursus maritimus*, with the probability of a reduction in population size by over 90% within 40 years set at 0.71. The IUCN have listed the polar bear as a vulnerable species ^7^, and understanding possible ways of adaptation to climate change in such an enigmatic animal at the genome level is key.

Environmental stress is reported to have a significant impact on genome structure, including the expression and mobilisation of transposable elements (TEs) ^8^. TEs are mobile genetic elements that can shape evolution through self-replication and insertion events in the genome, which in turn may change gene expression patterns ^9,10^. TEs inserted into or near a promoter region may alter the expression of genes and generate novel regulatory networks ^11^. TEs make up 38.1% of the genome in polar bears ^12^, which is similar to the giant panda *Ailuropoda melanoleuca* at 37%, and 36.1% of the dog *Canis familiaris* genome ^13^, and hence TEs may play an important part in the process of adaptation. Current data show that over 150,000 genome insertion mutations have been generated by TEs, driving variation in *ursine* and *tremarctine* bears ^12^. The aim of this study was to test if TE divergence between SEG and NEG polar bear populations may explain genetic differences and variation in gene expression between these sub- populations using a published transcriptome dataset ^14^. We included information on temperature variation into our analyses to test for a possible association between temperature and TE activity.

## Methods

### Meteorological analysis

We collected and analysed meteorological data on the climate in NEG and SEG from the Danish Meteorological Institute ^15^. SEG locations included: Aputiteeq, Tasiilaq, Mittarfik, Kulusuk, Ikermiit and Qaqortoq. NEG locations included: Ittoqqortoormiit, Daneborg, Port of Denmark and Station Nord (see Supplementary File 1 for latitude and longitude co-ordinates). Temperature data covering the highest, and lowest recorded temperatures each year from 1958-2024 were included and the data was plotted with Tableau Public version 2024.1.

### RNA-seq data

Transcriptome data from blood samples of adult polar bears were downloaded from the European Nucleotide Archive under accession no. PRJNA669153 ^14^ (see Suppl. File 2 for sample accession numbers and metadata) and aligned to the published reference polar bear genome ASM1731132v1. As described in the original publication ^14^, RNA-sequencing (RNA-seq) was based on total RNA and haemoglobin depletion step by CRISPR-Cas9 was used during library preparation. The libraries were sequenced on Illumina HiSeqX to generate 151 bp paired end reads ^14^. We selected adult samples with an average coverage of 28.4 million aligned reads, and included samples from five female NEG and seven female SEG bears and two male NEG and three male SEG bears in our downstream analyses. The average number of mapped reads across our 17 samples was 71 million. NF-core RNA-Seq pipeline version 1.4.2 was used to analyse the RNA-seq data (https://nf-co.re/rnaseq/1.4.2/) ^16^. Genome and annotation files were used and accessed as follows: ASM1731132v1 (GCF_017311325.1) GCF_017311325.1_ASM1731132v1_genomic.fna, and annotation GCF_017311325.1_ASM1731132v1_genomic.gtf https://www.ncbi.nlm.nih.gov/datasets/genome/GCF_017311325.1/). ASM1731132v1 is the most recent genome assembly and was used here as the reference genome. All differential expression (DE) analyses are comparing intercepts? of NEG versus SEG population.

### Differential TE expression analysis

To identify locus-specific expression of TEs and DE TE species, we used *Telescope* v1.0.3, a software previously tested on blood sample data ^17^. In addition to differential expression of TEs, we used *RepeatMasker* ^18^ 4.1.1 and *RepeatModeler* 2.0.3 ^19^ for estimation of Kimura distances to assess evolutionary divergence of nucleotide substitutions, TE activity and TE age: TEs with Kimura distance <10 are younger and more recently active. For this pipeline, we merged the bam files of individual samples by treatment group, then indexed and converted to fasta format with *Samtools* ^20^. *RepeatMasker* was used as follows: *RepeatMasker -species 29073 -par 5 -s -a -nolow -no_is*, with fasta inputs split into 200 chunks for parallelisation. Aligned outputs were merged and calcDivergenceFromAlign.pl was used for analysis. Genome sizes in the converted fasta files were calculated using calcGenomesize.pl. This version of *RepeatMasker* utilised Dfam v3.2, with genome version UrsMar1.0 TE annotations. The RepeatMasker default library for TEs in polar bears utilises an UrsMar1.0 based annotation, therefore a custom library, “-lib”, was generated for the ASM1731132v1 reference genome assembly. To generate a custom library “- lib” for use with RepeatMasker with the GCF_017311325.1_ASM1731132v1_genomic.fna as reference genome for the ASM1731132v1 assembly. To create a *RepeatModeler* database*: BuildDatabase -name custom_repeats_db_ASM1731132v1 -engine ncbi “$GENOME_FILE”*. Followed by: *RepeatModeler -database custom_repeats_db_ASM1731132v1 -pa 40 -engine ncbi*. The output was used as the -lib input instead of species in the RepeatMasker pipeline. RepeatModeler 2.0.3, ncbi-blast 2.10.1 and rmblast 2.10.0 ^21^, were used to analyse TE divergence. To produce the TE gtf annotation file for Telescope pipeline, the Perl scripts from TETranscripts ^22^ were used as follows: *perl makeTEgtf.pl -c 1 -s 2 -e 3 -o 6 -t 4 ASM1731132v1_rmsk > ASM1731132v1_rmsk_TE.gtf.* To test for a significant shift in the age of Kimura substitution values, R scripts for general linear modelling were used with R packages *MASS* v7.3-60.0.1 ^23^ *pscl* v1.5.9 ^24^., with a negative binomial model (Count ∼ Condition + Div; where Count: the TE abundance at the Kimura substitution level; Condition: enrichment in SEG population; Div: the Kimura substation levels) showing the best fit due to overdispersion in the TE count data (See Suppl. Table 1 for model comparisons). To analyse genomic features (coding sequence CDS, exon, gene, lncRNA, mRNA, pseudogene, snRNA and transcript) that overlap with significantly DE TEs, the loci of TEs were extracted from the Telescope results and mapped against gtf and bed file annotations to find genes and genomic features that overlap with bedtools intersect ^25^. All significantly DE TEs were tested for overlap with genomic features, and significantly enriched DE genes overlapping with DE TEs in the SEG population were used for GO term analysis.

### Differential gene expression analysis

We used *DESeq2* ^26^ in *RStudio* v 1.4.1717 with R v4.1.1 ^27^ to run a differential expression analyses between SEG and NEG bears. To account for spurious reads, any gene or TE alignment with less than ten reads was discarded before running *DESeq2*. The basic model in *DESeq2* to generate a dds object was *design = ∼ sex + population* (population referring to NEG or SEG) to account for potential sex differences in expression levels. To test for a temperature effect on TE expression, the average temperature over the year preceding blood sample collection recorded in the town nearest to the bear’s location was included in the analyses (mean temperature over 2016, bears sampled in 2017) in an extended model: *design = ∼ sex + temperature + population.* Results were extracted as follows: *res_temperature <- results(dds, name=”temperature”)*. Because location and temperature were colinear, the additive model was used without an interaction term to avoid redundancy. The significance threshold for DE genes and TEs was log_2_ fold change > 1 and a *p*_adj_ < 0.05. For GO term analyses, significantly we used *ShinyGO* v0.77 ^28^ on DE genes with a background list of all genes found to be expressed in this study, and figures were generated using custom Python scripts.

### Software and analysis

All R scripts and Python scripts used to generate figures can be found at: https://github.com/alicegodden/polarbear/. The following packages were used in the analysis and presentation of the data in this project: *DESeq2* 1.34.1 ^26^, *Tidyverse* 2.0.0 ^29^, *EnhancedVolcano* 1.12.0 ^30^, *ggplot2* 3.4.2 ^31^, *Readr* 2.1.4 ^32^, *tidyr* v1.3.0 ^33^, *viridis* v0.6.4 ^34^, *reshape* ^35^, *hrbrthemes* v0.8.7, ^36^ and *gridExtra* v2.3, ^37^, *MASS* v7.3-60.0.1 ^23^ *pscl* v1.5.9 ^24^.

## Results

### Meteorological analysis

The meteorological observation data from Danish Meteorological Institute (DMI), revealed vast variation in average lowest temperature measured from 1958-2024 between NEG and SEG (Fig. 1A), as denoted by the 95^th^ percentile size of dots representing more variation. The variance in observed lowest average temperatures was higher in SEG compared to NEG (Fig. 1B). Tasiilaq (SEG) showed the highest reported temperature, and Station Nord (NEG) showed the lowest recorded temperatures (Fig. 1A).

**Figure 1.**
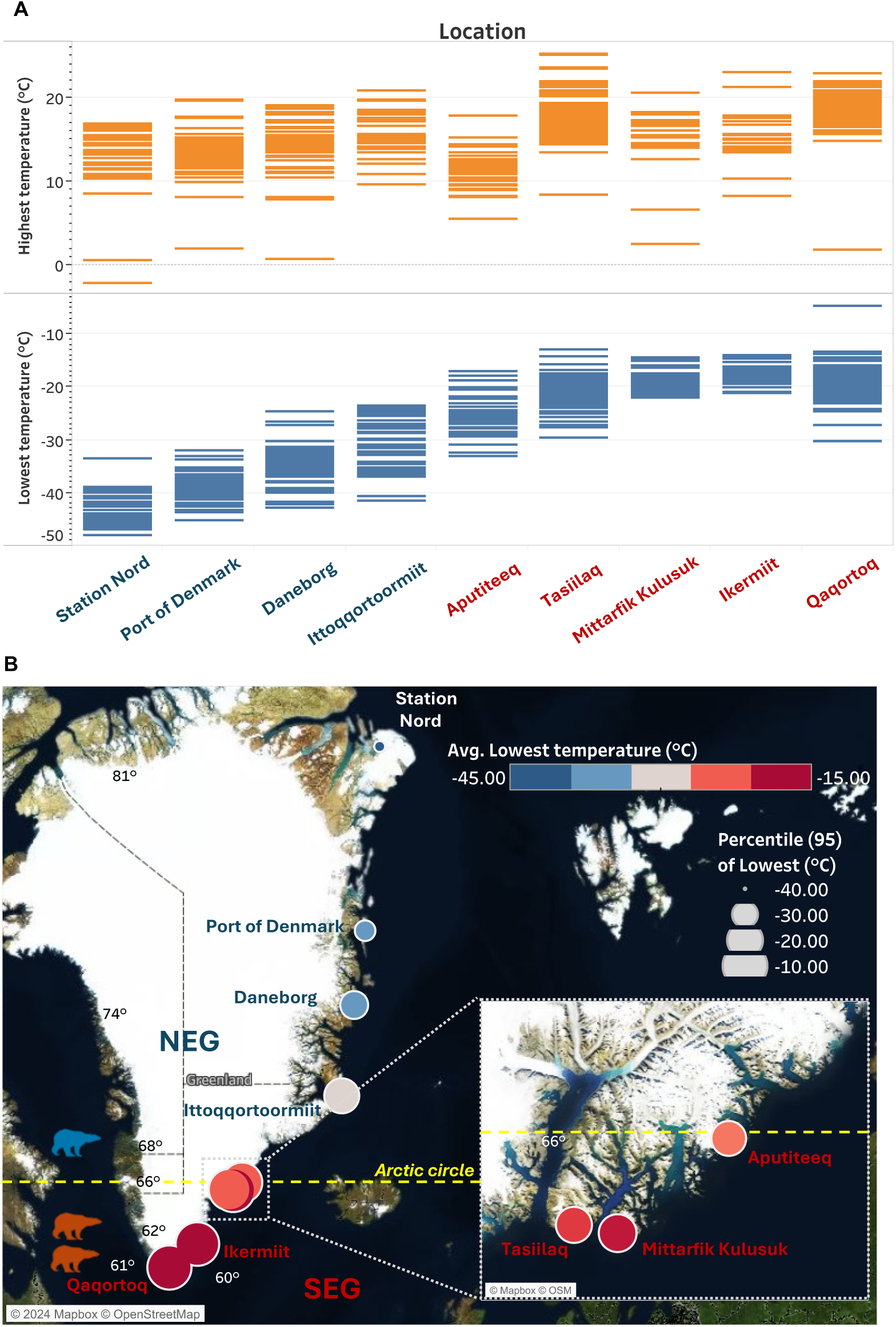
Meteorological observation data of temperature from the DMI. **(A)** Highest and lowest observed temperatures from 1958-2024, listed from Northernmost to most Southern. **(B)** Colour of points indicates the average lowest observed temperature, with the size of point showing the variance at the 95^th^ percentile. SEG locations included: Aputiteeq, Tasiilaq, Mittarfik, Kulusuk, Ikermit and Qaqortoq. NEG locations included: Ittoqqortoormiit, Daneborg, Port of Denmark and Station Nord. In this study north of 64° was considered NEG. The bear icons are indicative of the latitude where they were sampled, red meaning SEG and blue NEG. The Arctic circle is denoted by a dashed yellow line at 66°.

### Differential TE expression analysis

We initially identified 179 significantly DE TEs (*p_adj_* ≤ 0.05) between SEG and NEG bears when including both sex and population into the model (Fig. Suppl. Fig. 2B). A Principal Components Analysis showed modest clustering of samples based on population (Suppl. Fig. 2A). Most of the significantly DE TE species belongs to the LINE family (Suppl. Fig. 2B-C). The most significantly downregulated TE was DNA TE hAT-Charlie_dup1276 from the Charlie 8 subfamily. The most significantly enriched TE was LINE TE L1_dup3552 from the L1 Carn1 subfamily (Suppl. Fig. 2B).

When adding temperature as a covariate into our analyses, samples clustered based on temperature (Fig. 2B), and we identified 1,534 significantly DE TEs (Fig. 2A,C). The most significantly enriched TE was L2b LINE L2_dup45185, and the most significantly downregulated TE was L2a LINE/L2 dup_44480 (Fig. 2A). The most abundant TE class were again LINEs (Fig. 2C). To find genomic regions where TE activity may originate from, the genomic positions of significantly DE TE species were plotted (Fig. 2D), revealing some clustering of DE TEs on chromosomes NW_024423453.1, NW_024424119.1 and NW_02442563.1. No significantly DE TE species were identified on the Y chromosome (NW_024423336.1), but a few LINE TE species were located on the X chromosome (NW_024423319.1).

**Figure 2.**
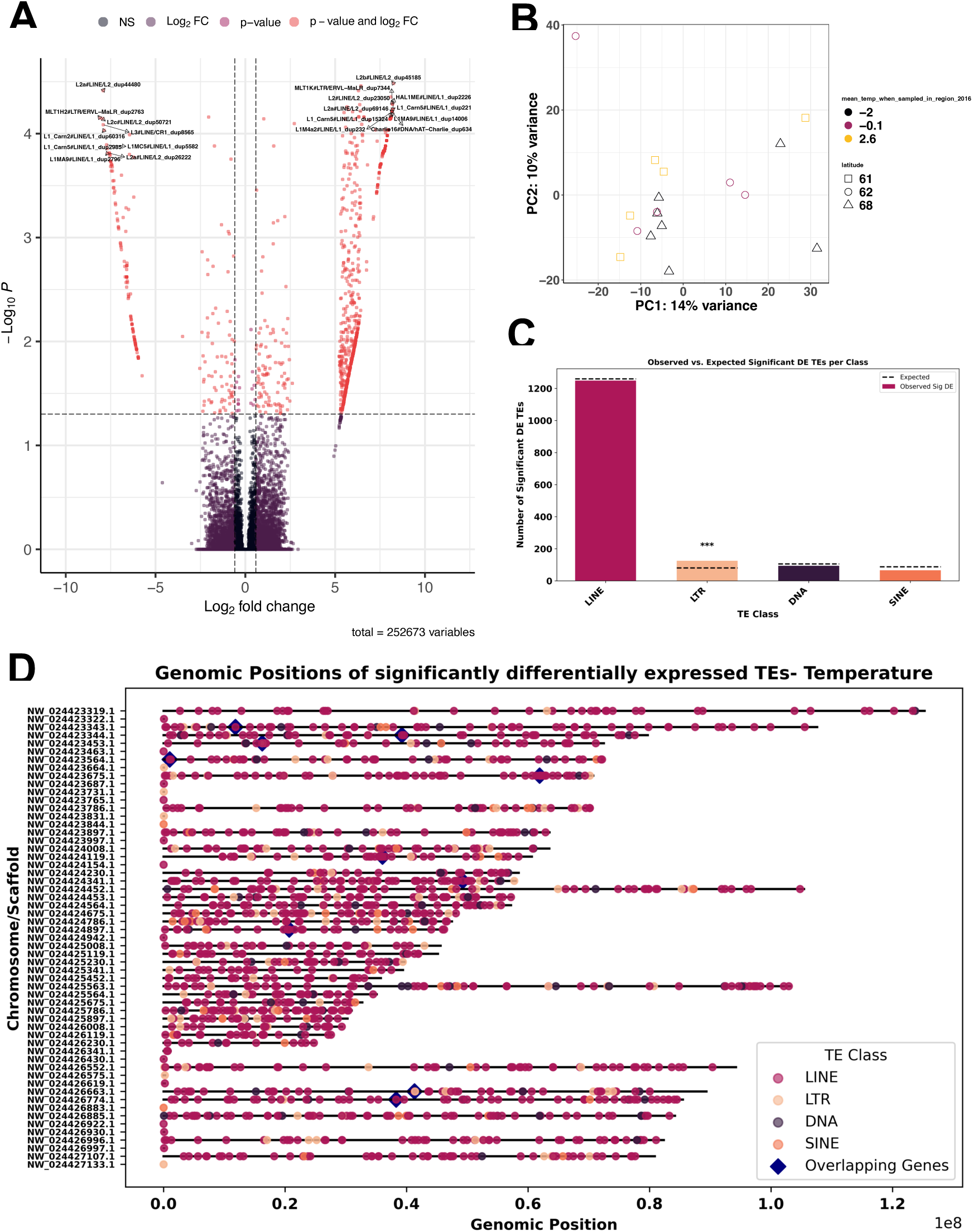
Comparison of expression and activity of TEs in NEG versus SEG bear taking temperature into account, profiling impact in SEG population. **(A)** Differential expression analysis from DESeq2 analysis of TE species identified 173 significantly differentially expressed TEs (*padj* ≤ 0.05) by the Telescope pipeline, with p cut off for significance *p*adj ≥ 0.05 on y axis and log2 fold change more than 0.5 (1). **(B)** Principal Component Analysis of in SEG and NEG polar bear populations including temperature and location. **(C)** Count data of significantly differentially expressed TEs at the family level as tested by hypergeometric testing (*** = *p ≤0.001*). **(D)** Genomic position of DE TE species and co-located DEGs.

Age estimations of DE TEs showed significant divergence in TEs between the SEG and NEG bear populations (Fig. 3), with both populations displaying divergence most strongly in LINE family TEs, followed by DNA and LTR TEs. In the SEG population, we saw an overall increase in TE divergence and a notable peak of younger divergent LINE TEs (Fig. 3). To examine if the significantly DE TEs are overlapping with genic regions, we mapped the TE loci against the bear genome. We found a significant enrichment of TEs in genic regions across the whole genome and not restricted to DE genes (Suppl. Fig. 1A). We profiled the TE-overlapping genes enriched in the SEG population with GO enrichment and found several significantly enriched KEGG pathways, biological processes and molecular functions in SEG population. Of relevance to a changed environment for the SEG bears, there was enrichment in KEGG pathways (Suppl. Fig. 1B) for *Foxo* signalling, metabolism changes (ovarian steroidogenesis, Thyroid hormone signalling) and immune response pathways including (human papillomavirus infection, chemokine signalling and cancer pathways). For biological processes there was significant enrichment of cardiac processes (SA node cell signalling pathways), metabolic processes (phosphorous metabolic processes), and a range of developmental processes including nervous system development (Suppl. Fig. 1C) and binding (Suppl. Fig. 1D).

**Figure 3.**
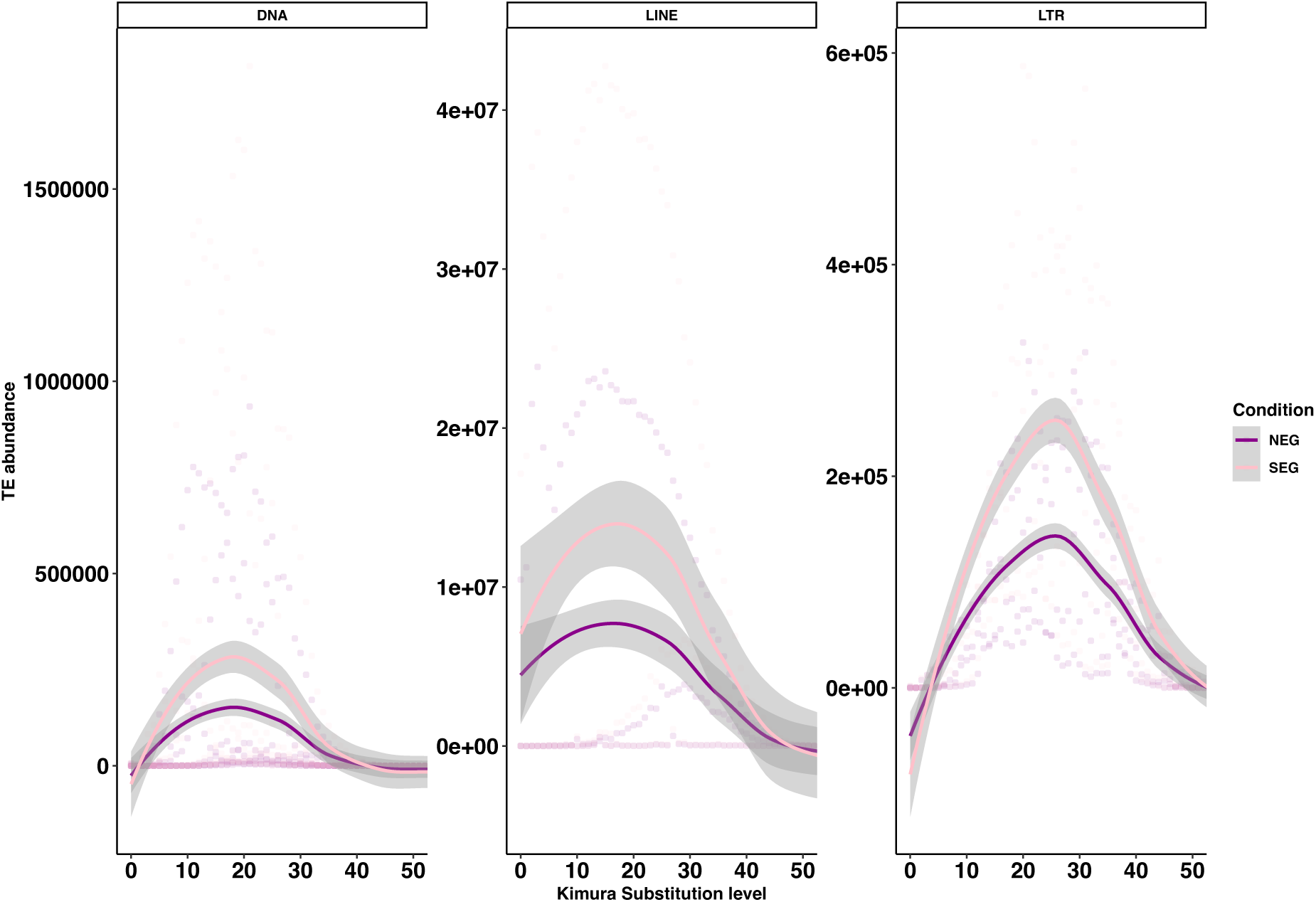
Repeat landscape of TE age and activity in NEG v SEG populations. TE age and divergence estimation by Kimura substitution analysis with RepeatMasker. The estimated divergence for SEG bears was greater than that in NEG bears. Analysis of negative binomial testing with general linear modelling results to assay the model *Count ∼ Condition + Div*, for a shift in TE age with negative binomial modelling showed significant increase in younger TEs in SEG populations at *p* ≤ 0.001 for all TE families. See Suppl. Table 1 for full statistical results.

### Differential gene expression analysis

When running the model including *sex + temperature + population* as fixed variables, clustering analysis revealed grouping of samples on PC1 with up to 29% variance explained by temperature and latitude of sampling and 15% in expression (Fig. 4). From this DE analysis, we identified 27 significantly DE genes, with LOC103677657 most downregulated and LOC103668192 most enriched (Fig. 4B). We found no overlap in position of significantly DE genes and TEs, but when using sex + population covariates, but addition of temperature covariate for ASM1731132v1 revealed 10 overlapping genes with TEs (Fig. 4D). These genes were: LOC121104823, LOC103667593, LOC103678552, LOC103677657, LOC121100717, LOC121101012, BBS12 (ENSUMAG00000004608- Bardet-Biedl syndrome 12), LOC121104992, SHANK1 and LOC103660844.

**Figure 4.**
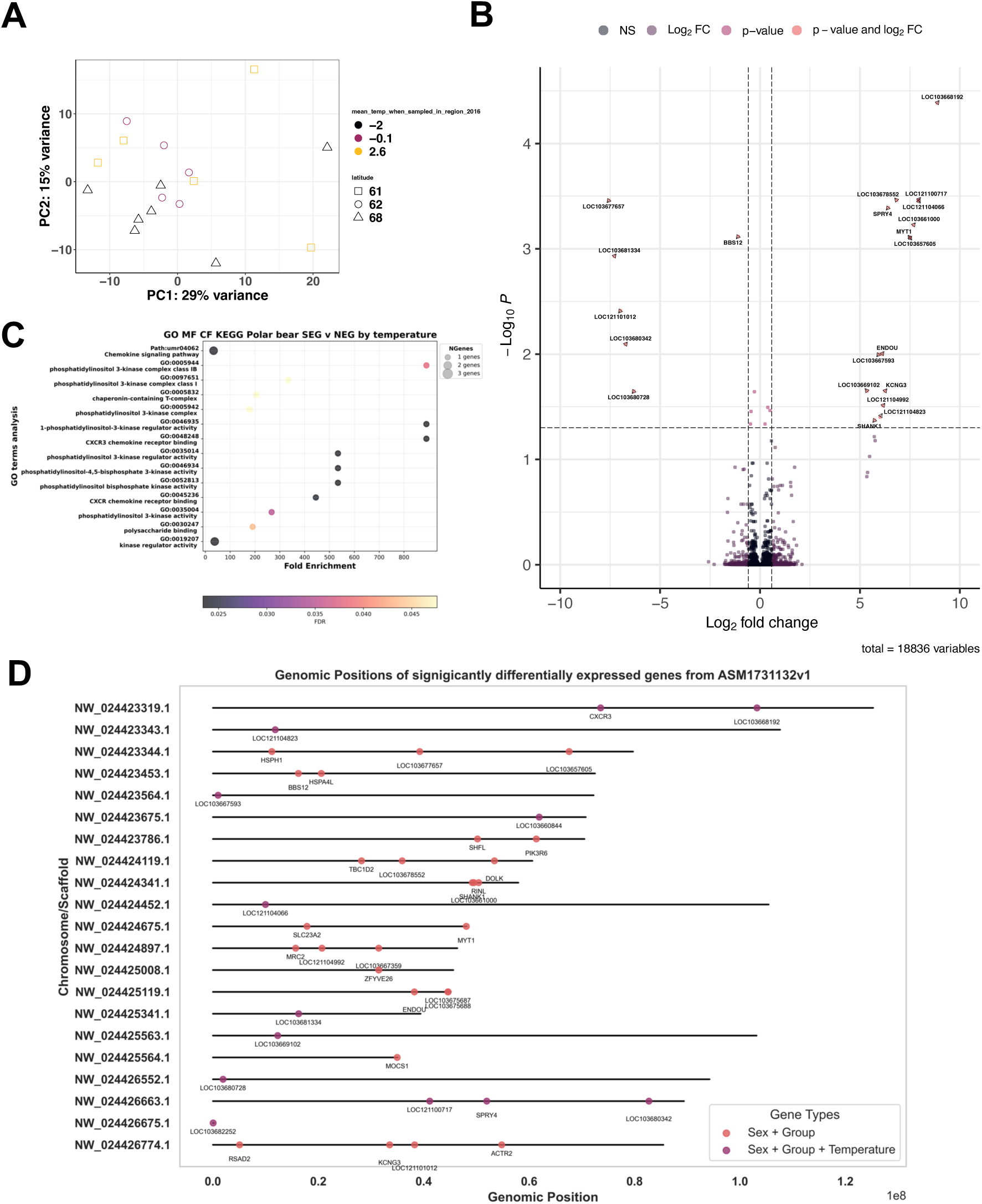
Analysis of differential gene expression to assay the main impact of temperature on the genomes in DESeq2 outputs looking at changes in SEG population. **(A)** Principal Component Analysis of differentially expressed genes in NEG and SEG polar bears taking temperature and sex into account shows clustering according to geographical location. **(B)** Volcano plot of differentially expressed genes, with p cut off for significance *p*adj ≥ 0.05 on y axis and log2 fold change more than 0.5 (1), showing 27 significantly DE genes. **(C)** GO terms analysis of significantly differentially expressed genes between SEG and NEG bears. **(D)** Loci of significantly differentially expressed genes with and without temperature covariate in *DESeq2* analysis.

Differential gene expression analysis for the model with *sex + population* as fixed variables revealed 13 significantly DE genes (Suppl. Fig. 3B). RSAD2 was most significantly downregulated and LOC10367587 was most significantly enriched in SEG compared to NEG bears. Of note, HSPH1 and HSPA4L were significantly enriched in SEG bears (Suppl. Fig. 3B). GO terms analyses of biological processes, molecular function, and cellular function for the significantly DE genes overlapping with DE TEs when aligned to ASM1731132v1 showed enrichment for the following terms: internal cellular dynamics and movements: chromosome movement towards spindle pole, PRC1 complex, organic acid:sodium symporter activity, negative regulation of protein localization to nucleus, and immune responses: negative regulation of viral genome replication, defence response to virus, defence response to symbiont, (Suppl. Fig. 3C).

## Discussion

The average lowest temperatures were highest in SEG towns in Greenland (Fig.1) suggesting bears in this region to be exposed to a significantly warmer climate, which can trigger an increase in TE activity. In accordance with this hypothesis, we found significantly DE TEs and genes between SEG and NEG bears (Figs. 2-4). The most abundant DE TEs between the two populations were LINE family TEs, and LTR TEs were significantly enriched in the SEG bears (Fig. 2C). We also found that the loci of significantly DE TEs overlapped with genic regions of the bear genome (Suppl. Fig. 1A). Kimura substitution analyses further showed an increased abundance of younger DNA, LINE and LTR TE s in SEG bears (Fig. 3). Significantly DE genes enriched in SEG polars included heat-shock protein genes HSPA4L and HSPH1 (Suppl. Fig. 3B). Taken together, our results suggest that TE activity in SEG bears is increased and may contribute to the differential expression of genes between the two populations. Below, we discuss the implications of our findings in a more general context and how they may help understanding adaptation in wild animal populations more generally.

### Meteorological analysis

Polar bears are dependent on sea ice, and the warmer and more variable SEG climate (Fig. 1A), is forecast to be the norm for all polar bears by the end of the 21^st^ century ^14^. The warmest temperatures in this study were observed in Tasiilaq (Fig. 1B), ranging between -30°C and +26°C, which is characterised by a deep fjord and mountainous habitat with some drifting sea pack ice as evidence of how temperature is isolating the SEG bears ^6,38^.

### Genetic divergence between SEG and NEG polar bears

When comparing the genetic divergence between SEG and NEG bears, a previous study reported an *F*_ST_ = 0.059 and concluded that SEG bears are more genetically diverged than other polar bear populations in Greenland ^14^. This genetic isolation is likely due to the limited gene flow caused by the East Greenland coastal current ^14^, restricting bear movements and thus promoting genetic drift over several hundred years of isolation. Despite this divergence, a previous study using the same dataset reported no significantly DE genes, in contrast to our findings of 27 DE genes. This discrepancy may be attributed to differences in model design, such as the inclusion of intercept terms and the specific covariates considered during analysis. The previous analyses were based on data from 16 adult samples, whereas we used 17 adult samples and used more stringent DESeq2 analysis, filtering out all genes with less than 10 reads, compared to the 1 read threshold previously analysed. The PCA showed similar clustering patterns between our study and the previous study.

### Differential TE expression analysis

The main clusters of DE TEs across all samples identified in our analyses are LINEs, and they were also the most abundant and significantly enriched in SEG bears (Fig. 2). LINEs and SINEs are the most abundant family of TEs in mammalian genomes in general, with limited LTR and DNA TE retention ^11^. The DNA TEs detected in this RNA-seq analysis are likely being detected through read-through of genes, this is unsurprising given the significant number of TEs in this study that overlap with genic and coding regions of the genome (Suppl. Fig. 1A). LINE and SINE TEs are also the most mobile and active TEs in bear genomes, and are most active in brown bears, followed by polar bears ^12^. To see if climate was impacting their expression, we included impact of temperature in our analysis. This identified 1,534 significantly DE TEs (Fig. 2A, C). This linking of temperature with genome mobility in polar bear aligns with similar studies in model systems including *Arabidopsis* ^39^, *C. elegans* ^40^ and *Drosophila* ^41^, where environmental stress was reported to increase TE mobility. Future work could include the use of DNA-sequencing (DNA-seq) and small RNA-seq data to analyse for TE insertion events and differential expression analysis of piRNAs.

Our finding of significant DE TEs (Fig.3) is in line with our idea of a putative role for TEs in the adaptation of SEG polar bears and offers new avenues to understanding rapid adaptation to changing environmental conditions. Divergent TEs between populations have also been described in great pandas, where LINE and SINE TEs were generally younger in age, indicative of recent mobilisation events ^13^. We found a high abundance of LINE TEs in both populations, potentially highlighting recent genome expansion of RNA class 1 TEs (Fig. 2C). LINE have been previously shown to be important in the evolution of regulatory elements in the mammalian genome ^42^. This is interesting as we observed a peak of younger, divergent LINEs in the SEG population of polar bears (Fig. 3). Older TEs included a modest amount of DNA TEs and LTRs in older genome expansion events, although these divergent events constitute a smaller fraction of the genome. This expansion of young LINEs, SINEs and LTRs was statistically significant (Fig. 3). Nevertheless, further DNA-sequencing analysis are needed to proof integration into the genome. Overall, the observed significant enrichment of young TEs in SEG bears could be in response to their challenging habitat and climate, but we need further evidence to fully appreciate their integration back into the genome.

To test if TEs are being enriched in expression due to their localisation within a gene or associated promoter region, we mapped the loci of the significantly DE TEs loci against the reference genome and tested for overlap with genic regions. Due to limitations in the polar bear genome assemblies, we could not perform intronic mapping, which is a technical as limitation TEs are expected to fall into intronic regions ^43^. However, we did manage to map TE loci to a significant number of genes as well as to lncRNAs and pseudogenes (Suppl. Fig. 1A). We observed significant enrichment of KEGG pathways and GO terms associated to these overlapping genes (Suppl. Fig. 1). One of the key pathways, *Foxo* signalling, plays an important role in adaptation to environmental stress and in ageing ^44,45^. The significant enrichment of *Foxo* signalling could be associated with increased TE activity in response to environmentally induced DNA damage and repair ^45^. The KEGG pathway *ovarian steroidogenesis* was also enriched, genes in this process included: INSR and ALOX5. This is interesting as warmer environments can affect polar bear diet, and if polar bears have lower fat reserves they are more likely to have a failed pregnancy, with data previously showing a 28% reduction in the number of pregnant female bears due to warmer environments ^46^. Despite this, the upregulation of ovarian steroidogenesis and lipid binding may be a way to compensate for hormonal imbalances caused by temperature rise and nutritional deficits in the SEG population ^47^.

Increased TE activity has been shown to lead to immune responses in the host due to detection of DNA damage and subsequent inflammatory responses ^48^. This may explain the enrichment of processes for immune signalling pathways associated with significantly DE genes (Fig. 4B-C), and the genes that overlap with loci of significantly DE TEs (Suppl. Fig. 1). We estimate that the recent spike in TE activity, as demonstrated by the significant peak of younger TEs (Fig. 3) could be representative of at least a few hundred years, this would be enough to cover several generations of polar bears and is a long time to allow for introduction of new genetic variability due to TE activity to favour rapid adaptation ^8,49^.

While our study is assessing TE activity in a somatic tissue (blood), adaptation would rely on germline TE activity. Some TEs are tissue specific, and their activity is generally less tightly regulated in somatic tissues ^50^, providing a direct profile of a response to environmental stress and possible resulting genetic divergence that may in time lead to genomic adaptation ^8^. Nevertheless, the finding of differential TEs in a somatic tissue and the age difference in TEs between the two populations, suggests an overall difference in TE activity across several generations of bears in these two populations, possibly in response to differential environmental factors.

### Differential gene expression analysis

The evolutionary origins of bears as a taxonomic group are complicated, and they are thought to have evolved over the past 5 million years in harsh and rapidly changing environments ^51^. To better understand polar bear genomes, we therefore profiled the differential gene expression of SEG versus NEG polar bears, whilst using *sex* as a covariate to account for differences in sample numbers and sex biases in gene expression. Given the differences in habitat over the recent decades of extreme warming and variable weather patterns in SEG ^6^, it is unsurprising that SEG bears are starting to show significant differences in gene expression compared to NEG bears.

### Linking gene and TE profiles in response to temperature

Our re-analyses of gene expression using a more conservative filtering steps of the data revealed a number of significantly DE genes between the two populations when taking sex and location into account, which contrasts with earlier results on the same dataset reporting no significant DE genes ^14^. Upregulated DE genes in SEG bears included RSAD2 (downregulated), ACTR2, HSPA4L and HSPH1 (enriched) (Suppl. Fig. 3B), where RSAD2 (also known as Viperin) is an evolutionarily conserved antiviral protein-coding gene ^52^. ACTR2 (actin-related protein 2) in house mice *Mus musculus* is involved in thrombosis, and mutations in the gene have been found to be linked to thrombo-suppression ^53^ whereas HSPA4L (Hsp70) and HSPH1 (Hsp110) are highly conserved and related families of proteins known as heat-shock proteins universal to all eukaryotic species so far, acting as molecular chaperones ^54^. This is interesting as other Hsp chaperones, including *Hsp90*, regulate protein folding, unfolding and degradation ^55^. *Hsp90* and *Hsp70* are also important in piRNA biogenesis and silencing of TEs, and function in elevated temperature environments ^56^. The most significant differential expression in TEs was observed in the analysis with the addition of temperature as a covariate (Fig. 2A). This is supported by evidence in *Drosophila* where TEs are activated in response to heat shock due to *Hsp70* and other chaperones having a role in the collapse of piRNA production, and thus further TE action ^57^. Combined, this is evidence that the host genome is potentially responding to the changes in environmental temperature.

When taking temperature as covariate into account (Fig. 4B), out of our 27 significantly DE genes, 10 of these gene loci overlapped with significantly DE TEs, these included: LOC121104823, LOC103678552, LOC121100717, and LOC121104992 (all enriched), LOC121101012 (downregulated) are all long non-coding RNAs (lncRNAs) (Fig. 4B, D). Long non-coding RNAs are highly abundant in mammals, with over 80% of transcribed products generating lncRNAs that have roles in chromatin organization and gene regulation ^58^. The accessibility of chromatin, and methylation is significant as this can lead to de-repression of TEs^59^.

The most significantly downregulated gene in the SEG population was LOC103677657, a pseudogene with unknown function. BBS12, an mRNA for Bardet-Biedl syndrome 12 protein, involved in chaperone protein complex assemblies, and is orthologous with human *BBS12,* was another downregulated in the SEG bears. Mutations or loss of BBS12 in humans leads to cilia abnormalities ^60^. Significantly enriched genes in SEG bears with reported functions included LOC103667593 which is a predicted mRNA for filaggrin-2-like protein linked to inflammatory skin diseases in human ^61^, and LOC121104992, a predicted mRNA for keratin-associated 4-8-like protein. SHANK1 was also significantly upregulated in SEG bears and is an mRNA for SH3 and multiple ankyrin repeat domains protein 1 and orthologous to human *SHANK1*. *SHANK1* encodes a scaffold protein and can inhibit the activation of integrins ^62^, which can lead to changes in cell signalling. This links to the GO term processes observed for phosphatidylinositol activity highlighting cellular activity and signalling (Fig. 4C). Other GO terms in response to the warmer SEG climate showed enrichment for terms related to immune responses including defence response to virus, and negative regulation of viral genome replication (Suppl. Fig. 3C). TE invasion is known to induce an immune response in mammals, activating the interferon and innate immune pathways ^63^. Polar bears in regions of reduced sea ice habitat are at higher risk of pathogen exposure ^64^, which may explain the changes in SEG bears. However, whether SEG bears are indeed exposed to higher pathogen loads needs further careful investigations.

## Conclusions

Our work revealed key significant differences in the genomes of SEG bears compared to the NEG bears, most strikingly in the mobile genome, potentially because of more variable and higher temperatures in SEG. The main differences we found were the significantly DE TEs and recent divergent genome expansion events exhibited by LINEs. Our study provides a basis for further work into TEs in a wider subset of polar bear subpopulations and highlights potential candidate TE species for further investigation as potential markers of environmental stress. Future work to revise and further annotate the *Ursus maritimus* genome, at all levels not just TEs will prove highly valuable in the efforts to preserve this vulnerable species through genomic analyses. Larger scale analysis across all other subpopulations of polar bear, and comparative analyses across different bear species in a range of climate zones may prove highly valuable for the planning of conservation efforts and species management.

## Abbreviations

DE: differentially expressed
DMI: Danish Meteorological Institute
DNA-seq: DNA sequencing
lncRNAs: Long non-coding RNAs
NEG: North-East Greenland
piRNA: piwi-interacting RNA
RNA-seq: RNA sequencing
SEG: South-East Greenland
TEs: Transposable elements

## Acknowledgements

We thank Dr. Oliver Tam (Hammel lab), for TE annotation file generation support. This study was funded by grants from the Natural Environment Research Council (NE/S011188/1) and the European Research Council (SELECTHAPLOID - 101001341) to SI.

## Declarations

Authors declare no conflict or competing interests.

## Author contributions

AMG conceived the idea and conducted the bioinformatic analyses, and wrote the manuscript. BR contributed to data analysis and conducted metereological analyses. SI supported manuscript preparation and funding.

## Data and code availability

RNA-seq data was deposited by ^14^ under ENA accession PRJNA669153. All R and python scripts used to analyse this data can be found in the following Github repository: https://github.com/alicegodden/polarbear/. Metadata used for the files accessed can be found in Supplementary file 2. The datasets supporting this article have been uploaded as part of the supplementary material in files 1-8. Supplementary files include all raw counts data from RNA-seq analyses, DESeq2 outputs for differential gene and TE expression with different models can be found here: https://github.com/alicegodden/polarbear/tree/main/supplementary_data.

## SUPPLEMENTARY FILES

All supplementary files can be found here: https://github.com/alicegodden/polarbear/tree/main/supplementary_data

**Supplementary File 1-** Meteorological data accessed from DMI **–** “Suppl. File. 1- Temperature_data - DMI.csv”

**Supplementary File 2**- Metadata of samples used for the RNA-seq analysis “Suppl. File. 2. bear_metadata_adult.csv”

**Supplementary File 3 –** Raw counts RNA-seq ASM1731132v1: “Suppl. File 3-Raw counts RNA-seq ASM1731132v1.tsv”

**Supplementary File 4 –** DESeq2 data RNA-seq ASM1731132v1: “Suppl. File 4-DESeq2 data RNA-seq ASM1731132v1.csv”

**Supplementary File 5 –** Telescope raw counts RNA-seq- TEs ASM1731132v1: “Suppl. File 5-Telescope raw counts RNA-seq- TEs ASM1731132v1.zip” compressed csv file.

**Supplementary File 6 –** Telescope DESeq2 data RNA-seq ASM1731132v1: “Suppl. File 6- Telescope DESeq2 data RNA-seq ASM1731132v1.csv”

**Supplementary File 7-** DESeq2 data RNA-seq ASM1731132v1 Temperature effect: “Suppl. File. 7- Temperature_effect_results_newgen_bear.csv”

**Supplementary File 8-** DESeq2 data RNA-seq ASM1731132v1 Temperature effect on TE expression: “Supp. File. 8- Temperature_effect_results_newgen_bear_TelescopeTEs.txt”

---

**Supplementary Figure 1.**
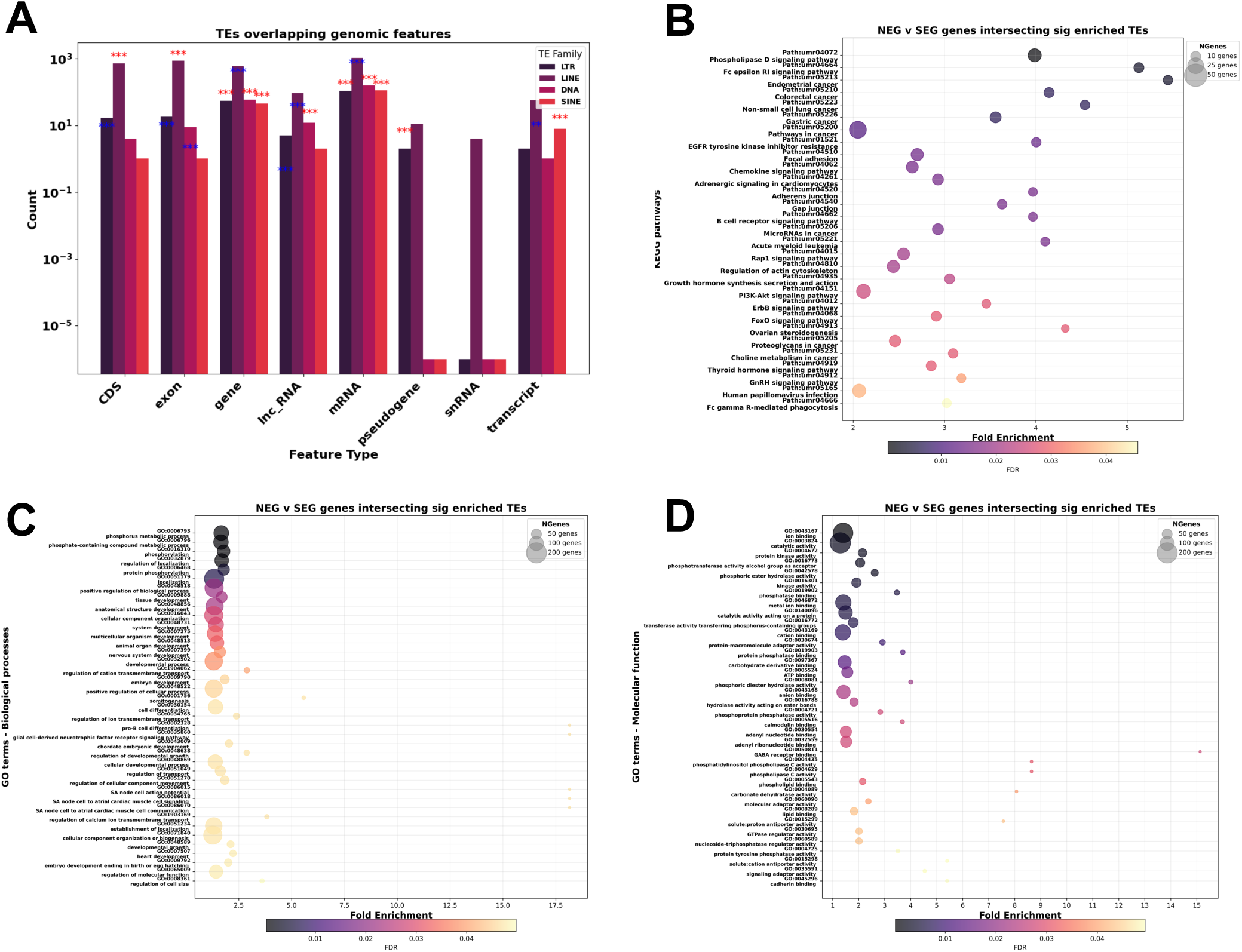
Significantly DE TE loci and their overlap with genomic features and genes in NEG v SEG populations. **(A)** Abundance of TEs overlapping genomic features, statistical enrichment analysed if this overlap was more significant than to be expected by chance (χ^2^ test of independence), *p* ≤ 0.001, as denoted by ***. SINEs were most significantly enriched in gene regions (χ² = 1036.94) and LINEs were most significantly depleted in transcript regions (χ² = 9.12). Stars in red show significant enrichment, stars in blue denote significant depletion. **(B)** KEGG pathways of overlapping genes. GO terms analysis of genes overlapping with significantly enriched TEs for **(C)** biological processes and **(D)** molecular functions.

**Supplementary Figure 2.**
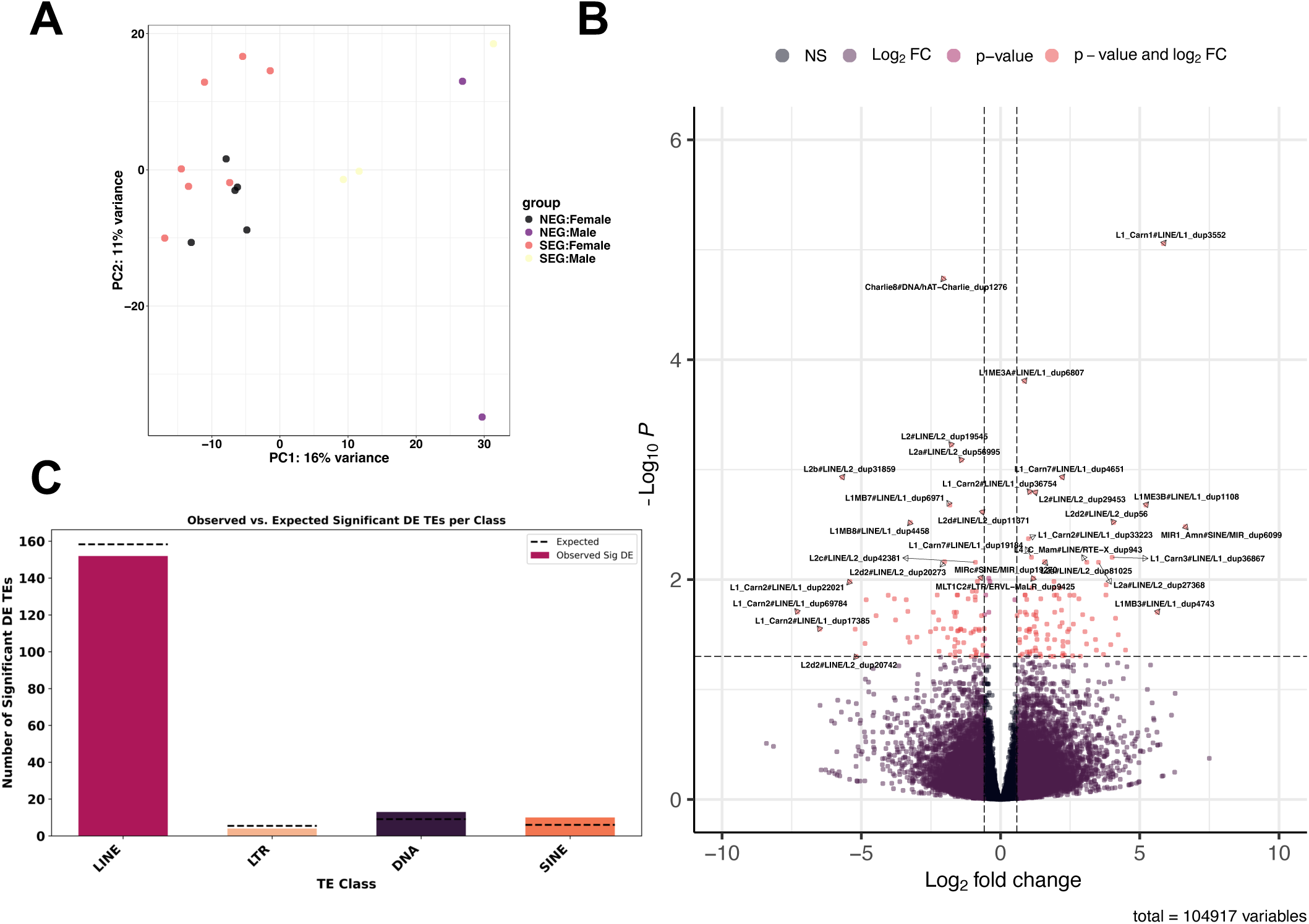
Comparison of expression and activity of TEs between SEG and NEG bears modelled on *sex + population* **(A)** PCA analysis of NEG v SEG bears show clustering based on geographical location of sample for TE expression. **(B)** Differential expression in DESeq2 analysis of TE species identified 173 significantly differentially expressed TEs (*padj* ≤ 0.05). **(C)** Count data of significantly differentially expressed TEs at the family level. **(D)** Genomic position of TE species that were significantly differentially expressed.

**Supplementary Figure 3.**
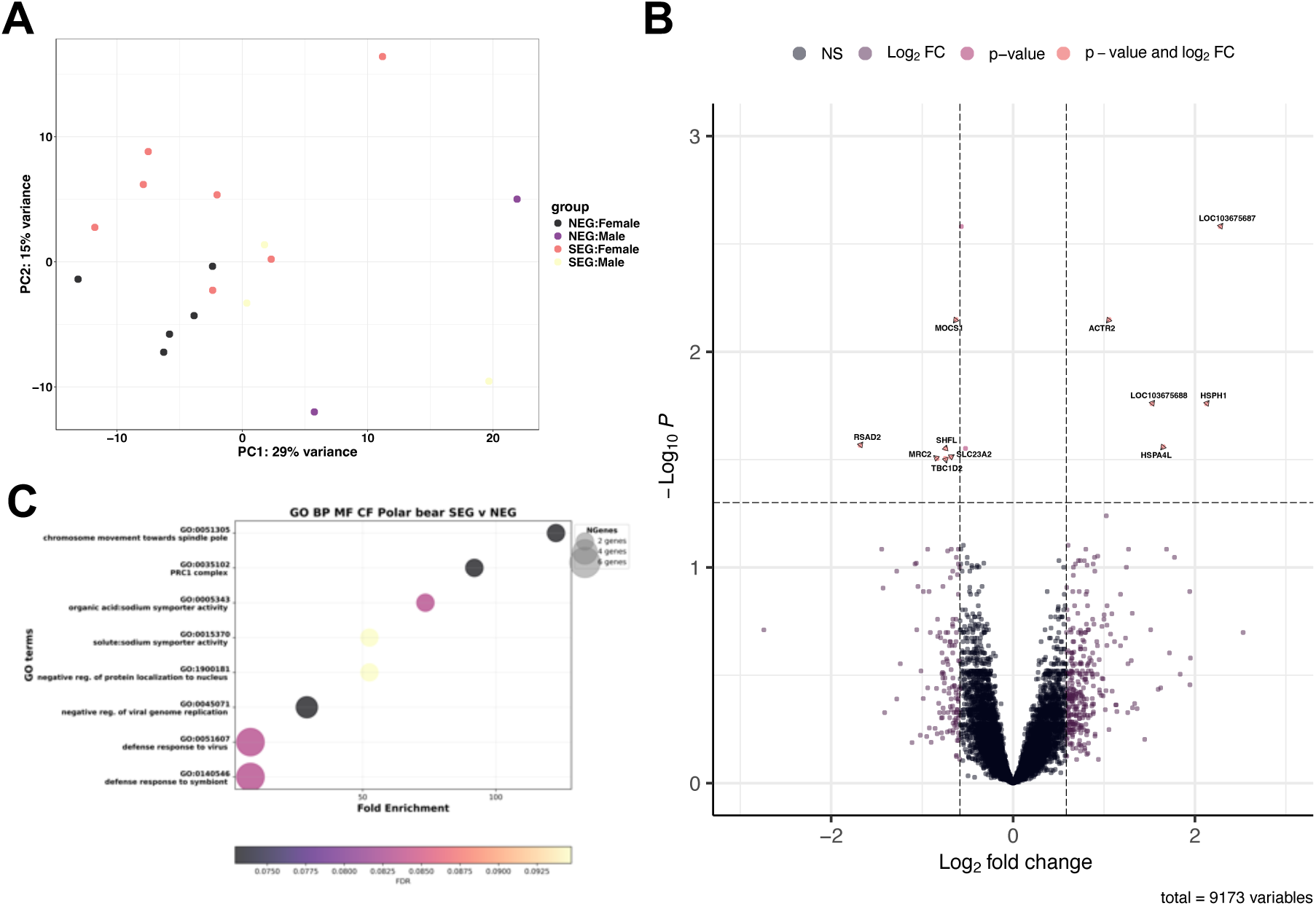
Differential expression analysis in *DESeq2* using RNA-sequencing data from samples in PRJNA669153 aligned to the reference genome ASM1731132v1 to compare NEG and SEG bears and observe impact on SEG population. **(A)** Principal Component Analysis shows variation in gene expression correlates to geographical location of the sample. **(B)** Volcano plot of the differentially expressed genes observed following DESeq2 analysis, with 13 significantly differentially expressed genes in total, with cut off for significance *p*adj ≥ 0.05 on y axis and log2 fold change more than 0.5 (1). **(C)** GO terms analysis, examining biological processes (BP), Molecular function (MF) and cellular function (CF). All genes expressed in the study were used as a background list, and those with *p* ≤ 0.1 processed for GO terms analysis with ShinyGO.

**Supplementary Table 1.** Statistical analysis of models to examine a shift in TE family age from Kimura Substitution analysis data from RepeatMasker outputs. General linear modelling results to assay the model *Count ∼ Condition + Div*, based on Akaike Information Criterion (AIC) and overdispersion values. Lower AIC values indicate better model fit. The Negative Binomial model is preferred over the Poisson model due to significant overdispersion in TE counts. The lower section of the table presents the results of a shift analysis, showing the estimated shift change, statistical significance (*p*-value), and the inferred shift direction in SEG polar bears Negative shift values and significant indicate that younger TEs are enriched in SEG regions.

| **TE_Family** | **Model_Type** | **AIC** |  | **Overdispersion** |
| --- | --- | --- | --- | --- |
| DNA | Poisson | 244884562 |  | NA |
| DNA | Negative Binomial | 244599505 |  | 457503.742 |
| LINE | Poisson | 4401719973 |  | NA |
| LINE | Negative Binomial | 4396475509 |  | 11897277.4 |
| LTR | Poisson | 66546483.4 |  | NA |
| LTR | Negative Binomial | 66503766.4 |  | 137545.593 |
| **TE_Family** | **Model_Type** | **Shift change** | **P value** | **Shift direction in SEG** |
| DNA | Negative binomial | -0.0461 | <0.001 / *** | Younger |
| LINE | Negative binomial | -0.0469 | <0.001/*** | Younger |
| LTR | Negative binomial | -0.0230 | <0.001/*** | Younger |

